# The expanded universe of prokaryotic Argonaute proteins

**DOI:** 10.1101/366930

**Authors:** Sergei Ryazansky, Andrey Kulbachinskiy, Alexei A. Aravin

## Abstract

The members of the ancient family of Argonaute (Ago) proteins are present in all domains of life. The common feature of Ago proteins is the ability to bind small nucleic acid guides and use them for sequence-specific recognition – and sometimes cleavage – of complementary targets. While eukaryotic Ago (eAgo) proteins are key players in RNA interference and related pathways, the properties and functions of these proteins in archaeal and bacterial species have just started to emerge. We undertook comprehensive exploration of prokaryotic Ago (pAgo) proteins in sequenced genomes and almost tripled the number of previously analyzed genes of this family. In comparison with eAgos, pAgos are highly diverse and have likely spread by horizontal gene transfer. Many pAgos contain divergent variants of the conserved domains involved in interactions with nucleic acids and in target cleavage, while having extra domains that are absent in eAgos, suggesting that they might have unusual specificities in the nucleic acid recognition and processing. Many pAgos, including catalytically inactive variants, are associated with putative nucleases, helicases and DNA binding proteins in the same gene or operon, suggesting that they are involved in DNA processing. The great diversity of pAgos revealed by our analysis opens new ways for exploration of their functions in host cells and their use as potential tools in genome editing.

## Introduction

Ago proteins play the key role in the RNA interference (RNAi) pathways in eukaryotes. All known eAgos bind small RNA molecules and use them as guides for sequence-specific recognition of long RNA targets. Upon recognition of the target, it can be cleaved by the intrinsic endonuclease activity of the Ago protein (Liu et al. 2004; Miyoshi 2005; Rivas et al. 2005; Rand et al. 2004). Alternatively, Ago proteins, especially the members of the family that lack nuclease activity, can recruit partner proteins to the target RNA leading to its degradation and/or repression of its translation (Djuranovic et al. 2012; Behm-Ansmant 2006; Rehwinkel et al. 2005). Recognition of nascent RNA by some eAgos can also lead to modification of the chromatin structure – DNA and histone methylation - of the target locus (Kuramochi-Miyagawa et al. 2008; Sienski et al. 2012; Verdel et al. 2004).

The proteins that belong to the Ago family are also present in the genomes of many bacterial and archaeal species (Makarova et al. 2009; Swarts et al. 2014b). Structural and biochemical studies of pAgos have provided key insights into the mechanisms of RNAi in eukaryotes: they revealed that Ago proteins directly bind short nucleic acid guides and can cleave complementary targets (Ma et al. 2005; Parker et al. 2004, 2005; Song et al. 2004; Wang et al. 2008a, 2008b, 2009; Yuan et al. 2005, 2006). The same studies revealed that pAgos can associate with short DNA guides and preferentially recognize DNA targets, in contrast to all known eAgos (Yuan et al. 2005; Wang et al. 2008a; Parker et al. 2009; Sheng et al. 2014; Kaya et al. 2016; Miyoshi et al. 2016; Doxzen and Doudna 2017; Willkomm et al. 2017; Liu et al. 2018). Despite these differences, solved structures of several pAgos and eAgos combined with their sequence alignments revealed a conserved domain organization of these proteins (reviewed in (Swarts et al. 2014b; Lisitskaya et al. 2018)). All eAgos and all except one pAgos that were experimentally characterized to date possess four domains that are organized in a bilobal structure, with N- and PAZ (PIWI-Argonaute-Zwille) domains forming one lobe and MID (Middle) and PIWI (P-element Induced Wimpy Testis) domains forming another lobe (Song et al. 2004; Yuan et al. 2005; Wang et al. 2008b; Elkayam et al. 2012; Nakanishi et al. 2012; Faehnle et al. 2013; Schirle et al. 2014; Matsumoto et al. 2016; Park et al. 2017). The nucleic acids are bound between the lobes; the MID and PAZ domains interact with the 5’- and 3’-ends of the small nucleic acid guide, and the PIWI domain contains an RNase H-like fold with a catalytic tetrad of conserved amino acid residues involved in the target cleavage (Wang et al. 2008b, 2009; Sheng et al. 2014; Kaya et al. 2016; Miyoshi et al. 2016; Willkomm et al. 2017; Liu et al. 2018).

Until now only a few pAgos were characterized by structural or biochemical approaches (see Fig. 1B). At the same time, earlier genomic studies revealed that up to 32% of the Archaea and 9% of the Eubacteria with sequenced genomes contain genes encoding proteins from the Ago superfamily and showed that the diversity of pAgos is far greater than that of eAgos (Makarova et al. 2009; Swarts et al. 2014b). Indeed, many pAgos contain substitutions of key catalytic residues in the PIWI domain (and are probably inactive), a large class of pAgos lacks the N- and PAZ domains (‘short pAgos’), while many pAgos contain additional domains absent in eAgos, either in the same protein or as a part of putative operons. Furthermore, previous studies of the genomic context of pAgo genes revealed that catalytically inactive pAgos are usually associated with several types of nucleases from the SIR2, TIR, PLD, Mrr (that include proteins with novel RecB-like and RecB2 domains) or Cas4 families (Makarova et al. 2009; Swarts et al. 2014b). Mrr and Cas4 are related to a highly divergent family of PD-(D/E)XK nucleases (Bujnicki and Rychlewski 2001; Kinch et al. 2005). Furthermore, MpAgo and TpAgo were found in association with Cas1 and Cas2 nucleases within the CRISPR loci (Kaya et al. 2016). The diversity of pAgos revealed by genomic studies together with few examples of biochemically characterized proteins suggested new cellular functions of these proteins (Olovnikov et al. 2013; Swarts et al. 2014a, 2015a; Zander et al. 2017). Beyond understanding functions of pAgo in their host cells, analysis of pAgos may yield new proteins that can be potentially harnessed for biotechnology, in particular as an alternative to the CRISPR/Cas genome editing tools (Hegge et al. 2017).

**Figure 1.**
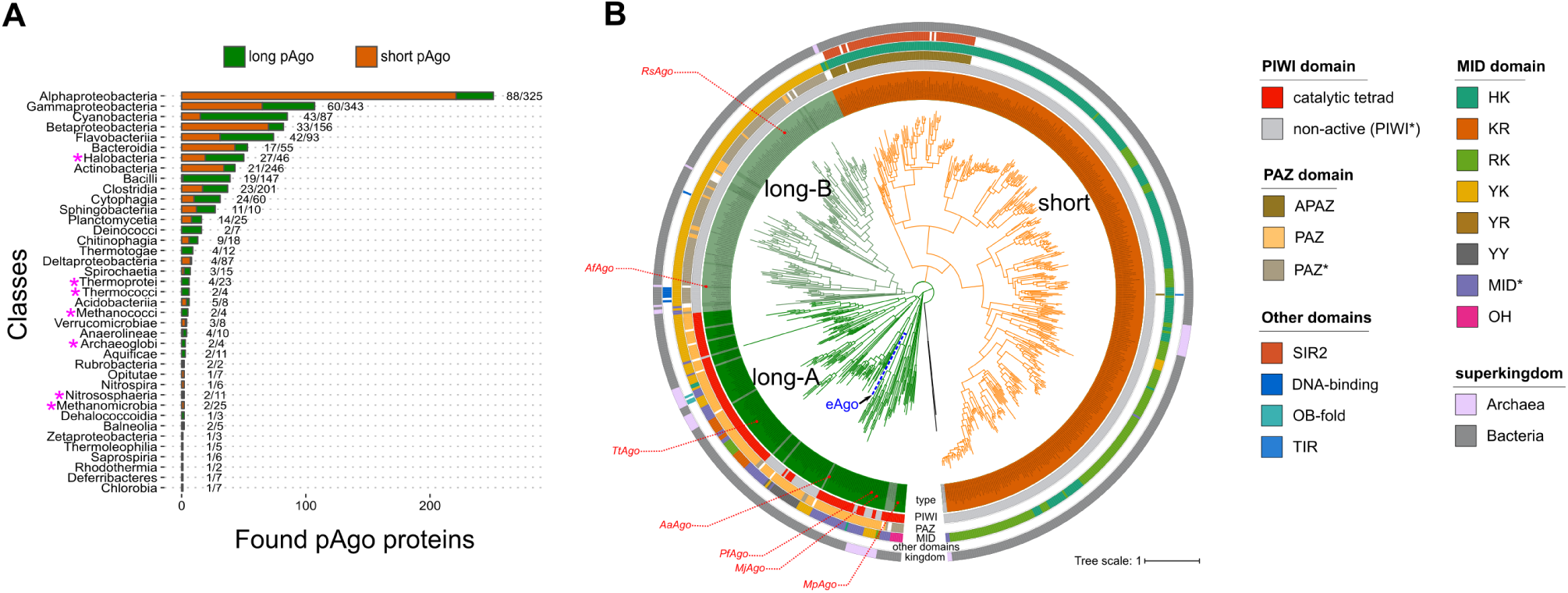
Phylogenetic analysis of pAgo proteins. **(A)**. The numbers of identified short and long pAgo proteins in the classes of Eubacteria and Archaea. Pink asterisks near the clade name indicate the Archaean classes. The numbers on the right of each bar depict the number of genera encoding pAgos *versus* the total number of genera with the complete or partially sequenced genomes in the corresponding class. (**B)**. The circular phylogenetic tree of the non-redundant set of pAgos constructed based on the multiple alignment of the MID-PIWI domains. The pAgo proteins were annotated as follows, from the inner to the outer circles: the type of protein, in which long-A pAgos are painted in green (truncated variants without the PAZ domain, light green), long-B pAgos are light-green (truncated variants without PAZ, green), and short pAgos are orange; the type of the PIWI domain, depending on the presence of the catalytic tetrad DEDX; the type of the 5’-end guide binding motif in the MID domain (the first two conserved residues are indicated); the presence and the type of the PAZ/APAZ domains in pAgos; the superkingdom to which the corresponding pAgo belongs to. Biochemically characterized pAgos (or the most similar redundant homologs for RsAgo and TtAgo) with resolved structures are highlighted in red. The scale bar represents the evolutionary rate calculated under the JTT+CAT evolutionary model.

Previous genomic studies identified a non-redundant set of 261 pAgos (Swarts et al. 2014b; Makarova et al. 2009). However, since their discovery the number of bacterial strains with sequenced genomes has more than tripled, and many previously sequenced genomes have been revised resulting in the removal of some pAgo genes from the databases. Therefore, we have undertaken comprehensive exploration of the diversity of pAgos using the latest version of the RefSeq protein database for bioinformatic search. The results of the search almost tripled the number of identified pAgo genes. Our analysis supports the previously reported separation of pAgos into short pAgos and long pAgos (which by themselves can be separated into two distinct clades), and reveal several subclasses with significant variations in the domain organization. These include variations in the MID and PAZ domains involved in guide binding and substitutions of catalytic residues in the PIWI domain. A large fraction of pAgos, including those that lack intrinsic endonuclease activity, are associated with putative nucleases and other DNA binding proteins. Overall, the great diversity of pAgo revealed by our analysis opens a way for exploration of their biochemical properties and cellular functions, as well as their use as potential tools in genome engineering.

## Results

### The expanded set of prokaryotic Argonaute proteins

To reveal the diversity of pAgos and to get new insights into their evolution we performed comprehensive analysis of proteins that belong to the Ago family in sequenced genomes of prokaryotic species. For this, we searched 116 million proteins fetched from the RefSeq NCBI protein database using known pAgos as queries by PSI-BLAST. We found 1010 pAgos encoded in 1385 completely and partially sequenced genomes of 1248 Eubacterial and Archaean strains (Fig. 1, Supplemental Table S1). In total, pAgos were found in 17% of Eubacterial and 25% of Archaean genera (Fig. 1A). The majority (1186, or 95%) of Archaean and Eubacterial strains encode only one pAgo gene (876 pAgos, or 86.7% of all proteins). However, the genomes of 57 strains encode two pAgo genes, four strains encode three pAgo genes, and one strain encodes four pAgos. pAgos are in general randomly distributed in different prokaryotic clades as the number of genera in each class that encode pAgos in their genomes correlates with the total number of genera in the class (Fig. 1A, Supplemental Fig. S1). The non-redundant set of pAgos with the level of similarity of less than 90% comprises 721 proteins, which is almost triple the number of previously identified pAgo proteins (Makarova et al. 2009; Swarts et al. 2014b).

As described below, the vast majority of pAgos have the MID and PIWI domains, but many proteins lack the N- and PAZ domains conserved in eAgos. Therefore, we used multiple alignments of the MID-PIWI domains to construct a phylogenetic tree of pAgos (Fig. 1B, Supplemental Fig. S2). The tree revealed three large clades of pAgos, while two proteins from thermophilic archaea *Thermoproteus uzoniensis* and *Vulcanisaeta moutnovskia* could not be unambiguously classified as belonging to any of these clades due to their high divergence. Following Makarova and co-authors (Makarova et al. 2009), we call one of these clades ‘short pAgos’ (Fig. 1B), as all proteins in this clade lack the N- and PAZ domains present in eAgos. The majority of pAgos that belong to the other two clades include the PAZ domain and we therefore designate them as long-A and long-B. The diversity of pAgos vastly exceeds the diversity of eAgos that were previously shown to form a single branch on the phylogenetic tree of long pAgos (as illustrated in Fig. 1B) (Makarova et al. 2009; Swarts et al. 2014b).

The three clades of pAgos can be found in both Archaean and Eubacterial species. Furthermore, the majority of prokaryotic classes that include large numbers of genera encode both short and long pAgos. However, some classes have a clear bias towards short or long pAgo types; thus, genomes of Alphaproteobacteria predominantly encode short pAgos, while Cyanobacteria predominantly encode long pAgos (Fig. 1A). pAgos that belong to different clades can be found in the same genome (for the strains that encode several proteins). For example, *Enhydrobacter aerosaccus ATCC 27094* encodes three short and one long-A pAgo; *Parvularcula bermudensis HTCC2503* and *Halomicronema hongdechloris C2206* encode two short and one long-B pAgos; *Methylomicrobium agile ATCC 35068* contains one short and one long-B pAgo. Generally, the phylogenetic tree of pAgos has little similarity with the phylogenetic tree of host species build using classic molecular markers such as rDNA, thus confirming previously published observations (Makarova et al. 2009; Swarts et al. 2014b) and suggesting that pAgos have mostly spread through horizontal gene transfer. Notably, a few pAgos that have been biochemically studied almost all belong to the long-A clade, while only two pAgos from the long-B clade and no proteins from the short pAgo clade have been characterized to date (indicated in Fig. 1B).

### Domain architecture of pAgos

To gain further insight into the structural diversity of pAgo, we analyzed their domain architecture. The exploration of multiple alignments of PIWI, MID and PAZ domains was used to identify the main structural and functional features of pAgos. Using InterProScan and CDD-batch programs and the Pfam and Superfamily databases we also searched for other domains in pAgo proteins. This analysis allowed us to propose a comprehensive classification of the pAgo domain organization (Fig. 2).

**Figure 2.**
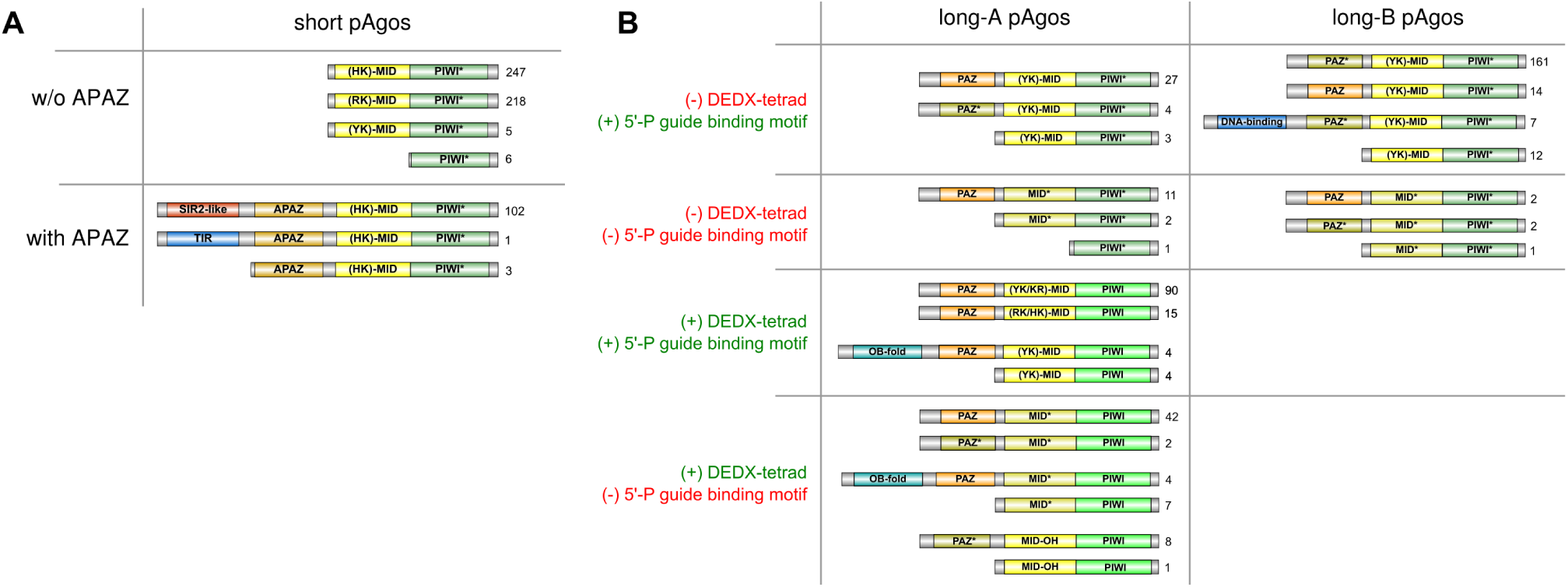
The diversity of domain architecture of pAgo proteins. Various types of domain organization of short (A) and long (B) pAgo proteins are schematically shown. The numbers on the right of each type are the frequency of its occurrence in the redundant set of proteins. Based on the presence of the DEDX tetrad in the PIWI domain and the putative motif 5’-P guide binding motif, all long pAgos can be separated into four types shown here. Green PIWI domains carry the active DEDX catalytic tetrad while turquoise PIWI* domains do not contain the canonical tetrad. Yellow MID domains contain different types of the 5’-end guide binding motif as indicated, while light-brown MID* domains carry substitutions of the crucial amino acid residues in this motif. Orange PAZ domains have full-sized pocket responsible for 3’-end guide binding, while light-brown PAZ* variants do not have the second subdomain. The OB-fold is the nucleic-binding domain SSF50249; the DNA-binding domain comprises Schlafen domain with AlbA_2 (PF04326) and lambda-repressor-like domain (SSF47413).

As justified by their name, we found that all members of the short pAgo clade lack the PAZ domain implicated in binding of the 3’-end of a guide molecule. The majority of short pAgos (470, or 81%) have only two domains, MID and PIWI. Six short pAgos (~1%) are further truncated and lack the MID domain suggesting that they may be unable to interact with a guide. In addition to the MID and PIWI domains, 106 short pAgos (18%) that all belong to a single monophyletic branch encompass the APAZ (‘analog of PAZ’) domain (Fig. 1B, Fig. 2A). The APAZ domain has no sequence similarity to PAZ and was previously described as a domain always associated with short pAgos either as a part of the same protein or present in the same operon with short pAgos (Makarova et al. 2009). Almost all (102 out of 106) APAZ-containing short pAgos also carry domains related to the SIR2 family (SIR2_1 subfamily) (Fig. 1B, Fig. 2A), which are considered to be *bona fide* nucleases (Makarova et al. 2009; Swarts et al. 2014b).

In contrast to short pAgos, the majority (378, or 90%) of pAgos that belong to the long pAgo clades have the domain architecture similar to eAgos, including PAZ, MID and PIWI domains (the N-domain that has much more diverse sequences was not included in our analysis). However, 31 pAgos that undoubtedly belong to long pAgos based on alignments of their MID-PIWI domains lack the PAZ domain, including previously characterized AfAgo from *Archaeoglobus fulgidus* (Parker et al. 2004; Ma et al. 2005; Parker et al. 2005). These truncated long pAgos are scattered at different positions on the phylogenetic tree suggesting multiple independent cases of the loss of this domain during evolution of long pAgos. One of these truncated proteins also lacks the MID domain. On the other hand, 15 long pAgos that have the PAZ domain also carry additional putative nucleic-acid-binding domains at their N-termini: 7 pAgos contain DNA-binding domains (Schlafen, PF04326, or lambda-repressor-like, SSF47413), and 8 pAgos contain a nucleic-acid binding domain with the OB-fold (SSF50249) (Fig. 1B, Fig. 2B).

Analysis of the diversity of MID, PIWI and PAZ domains revealed significant variations in their structures characteristic for different groups of pAgos (Fig. 1B, Fig. 2), as described in the next sections. In particular, subsets of proteins from both clades of long pAgos contain MID* domains with substitutions of key residues involved in interactions with the guide 5’-end; many long-A pAgos, all long-B and short pAgos contain PIWI* variants with substitutions of essential catalytic residues; and most long-B pAgos contain reduced variants of the PAZ* domain involved in the 3’-guide interactions (Fig. 2).

### Binding of nucleic acid guides in the MID domain

The vast majority (1001 or 99% of all proteins) of pAgos contain the MID domain that has been implicated in binding of the 5’-end of guide molecules. The resolved 3D structures of pAgos, including AfAgo (pAgo of *A. fulgidus*) (Parker et al. 2004; Ma et al. 2005; Parker et al. 2005, 2009), RsAgo (*Rhodobacter sphaeroides*) (Miyoshi et al. 2016; Liu et al. 2018), TtAgo (*Thermus thermophilus*) (Wang et al. 2008a, 2009; Sheng et al. 2014), and MjAgo (*Methanocaldococcus jannaschii*) (Willkomm et al. 2017), have shown that the MID domain anchors the phosphorylated 5’-end of the guide by a set of conserved amino acid residues (Fig. 3). Among them, four conserved residues (Y/R, K, Q and K in crystallized pAgos; *e.g*. Y463, K467, Q478 and K506 in RsAgo in Fig. 3C) as well as a bound divalent cation (Mg^2+^ or Mn^2+^) form a network of hydrogen bonds anchoring the 5’-phosphate, and also the third guide phopshate, into this basic pocket (Ma et al. 2005; Parker et al. 2005; Wang et al. 2008b; Miyoshi et al. 2016; Willkomm et al. 2017). Multiple sequence alignment of the MID domains shows that these residues are highly conserved in most pAgos (Fig. 3A and B). Accordingly, substitutions of these residues were shown to disrupt pAgo-guide interactions (Parker et al. 2004; Ma et al. 2005; Wang et al. 2008b; Miyoshi et al. 2016; Willkomm et al. 2017). Two additional semi-conserved residues (*e.g*. T and N in RsAgo; Fig. 3) contribute to binding of the phosphate group and the base of the second guide nucleotide, respectively (Fig. 3C). Together, these residues constitute a conserved six-amino-acid motif that is found in most pAgos (**YKQ**TN**K** consensus for long pAgos, with the most conserved residues shown in bold; indicated with red asterisks in Fig. 3A and B).

**Figure 3.**
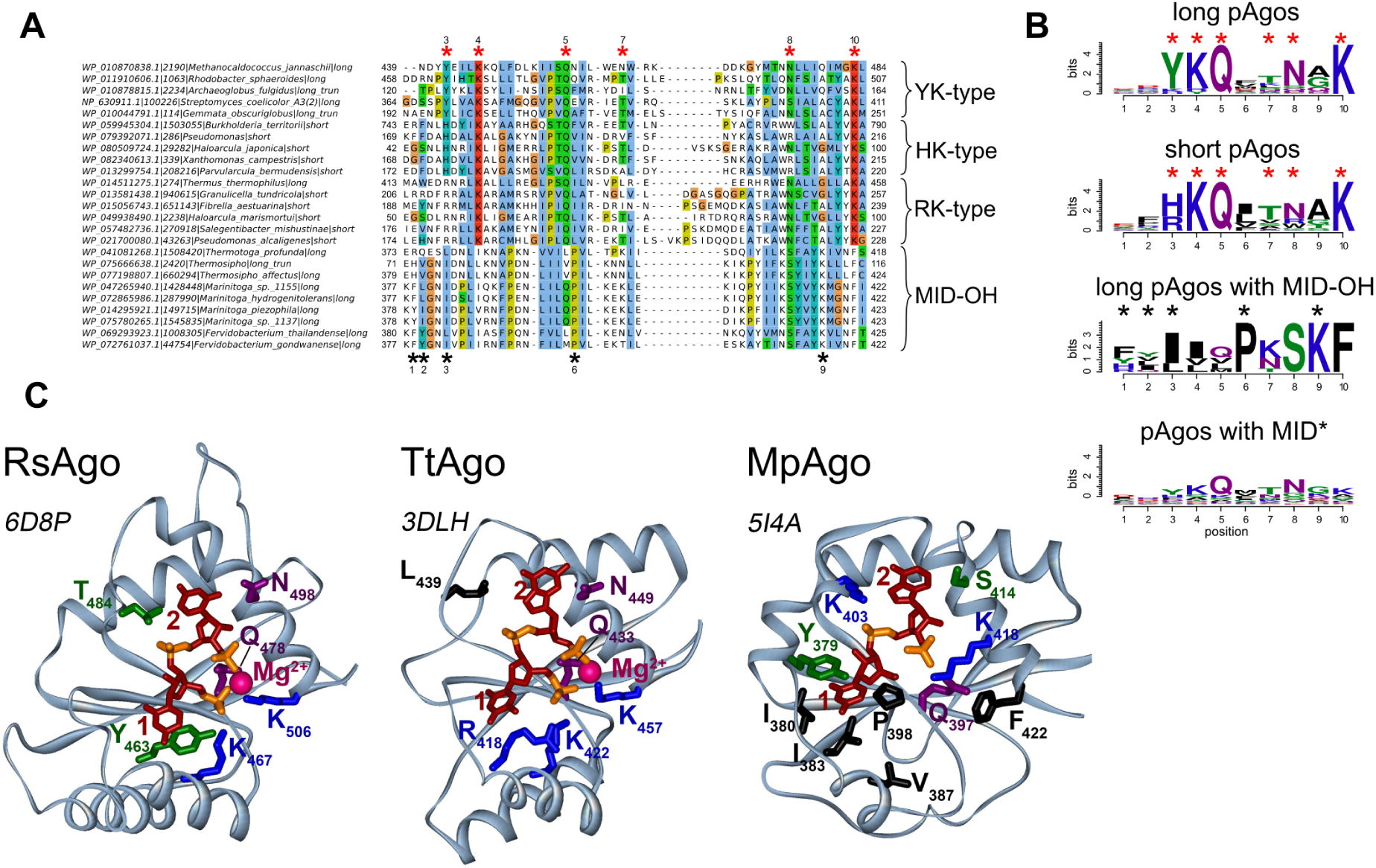
The structure of the 5’-end guide binding motifs in the MID domain. **(A)**. A fragment of the multiple alignment of MID domains of several typical pAgo proteins with different types of the 5’-end guide binding motifs. Red asterisks are the positions of amino acid residues that are involved in the binding of the 5’-P-end of a guide molecule. Black asterisks are the positions of amino acid residues involved in the binding of the 5’-OH-end of a guide in the MID-OH pAgos. MID domains with YK-type 5’-P-end binding motif belong to the long pAgos; MID domains of HK-type belong to the short pAgos; MID domains of the RK-type are characteristic to short pAgos but can also be found in some long pAgos. The multiple alignments were manually edited to bring them in conformity to the secondary structures shown in Fig. 3C. The species names, NCBI taxon_id, the types of pAgos (long, short and lomg truncated without the PAZ domain) and corresponding accession numbers are shown at the left of the alignment. **(B)**. The sequence logo of amino acid residues involved in the formation of the 5’-end binding pocket for long, short, MpAgo-like pAgos with MID-OH, and long pAgos with MID* domain variants. A combined logo motif including positions characteristic for canonical MID and MID-OH domains is shown. Red and black asterisks are amino acid positions specific for the 5’-P (MID) and 5’-OH (MID-OH) binding motifs, correspondingly. **(C)**. Structures of binary complexes of RsAgo, TtAgo, and MpAgo with guide molecules: 5’-P guide RNA for RsAgo, 5’-P guide DNA for TtAgo or 5’-OH guide RNA for MpAgo. For RsAgo and TtAgo, only residues from the conserved YKQTNK motif are indicated; for MpAgo, MID-OH-specific residues involved in interactions with the 5’-guide end are also shown (labeled with asterisks). PDB IDs are shown next to each structure.

Interestingly, we found amino acid variations at the first two positions of this motif, which might be important for the function of different groups of pAgos. The first two residues in the motif are usually YK in long pAgos; subsets of long-A pAgos contain KR, RK, HK or YY combinations, while long-B pAgos are almost exclusively YK variants. The aromatic ring of the first Y stacks on the first base at the 5’-end of a guide, thereby contributing to the binding of the first nucleotide within the MID pocket (Ma et al. 2005; Parker et al. 2005; Miyoshi et al. 2016; Willkomm et al. 2017; Liu et al. 2018). In contrast to tyrosine, R is positively charged and not aromatic and so unable to stack the first base of the guide, as exemplified in the structure of TtAgo (Fig. 3C) (Wang et al. 2008b; Sheng et al. 2014); the structure of any pAgo with histidine at the first position of the motif remains unknown.

The conserved residues implicated in binding of the 5’-phosphorylated guide end are present in the majority (80%) of long pAgos. However, 84 (20%) of long pAgos have at least one amino acid substitutions at the most conserved positions in the 5’-binding motif in their MID* domain; namely, they do not contain Y/H/R at the first position of the YKQTNK motif and/or K/R/Y at the second position, and/or Q at the third position, and/or K at the last position. A separate subgroup of these proteins (MID-OH), which form a tight clade on the phylogenetic tree, includes pAgo from *Marinitoga piezophila* (MpAgo) that was recently shown to bind guide molecules with 5’-OH instead of 5’-P ends (Fig. 1B) (Kaya et al. 2016). The multiple sequence alignment showed that this group includes 9 pAgos (as compared to 3 proteins identified in the previous study (Kaya et al. 2016) and that positions of amino acid residues involved in 5’-guide interactions in these proteins overlap with the positions of the 5’-P-end binding motif in other pAgos (Fig. 3A,B). However, in comparison with most pAgos, these residues form a more hydrophobic pocket for 5’-OH binding (Fig. 3C). In addition, the MID pocket of MpAgo does not contain bound Mg^2+^ ions. The absence of guide contacts with the Mg^2+^ ion is in part compensated by interactions between the second and third guide phosphates with lysine residues specific for this group of pAgos (K403 and K418 in MpAgo; Fig. 3B and C). While the first tyrosine of the YKQTNK motif that stacks with the first guide base in other pAgos is replaced with a hydrophobic residue in the MID-OH domain, its role is taken by a preceeding aromatic residue (Y or F) that stacks from the other side of the same base (Y379 in MpAgo; Fig. 3B and C).

Other 75 pAgos with non-canonical MID* variants, none of which was characterized to date, belong to several distinct branches on the phylogenetic tree of pAgos, and some of them are located close to the MID-OH pAgos (Fig. 1B). The alignment of these sequences showed that in general they share the same YKQTNK motif, although the conservation of individual positions is much lower than for canonical pAgos (see logo in Fig. 3B; alignment in Supplemental File 1). It remains to be established whether these proteins include any additional subgroups with noncanonical MID* domains of different specificities, as already revealed for MpAgo, or they just represent divergent variants of the same canonical motif.

In contrast to long pAgos, all short pAgos have the canonical 5’-P-end binding motif (Fig. 3B). Interestingly, almost all of them contain HK and RK residues at the first two positions of this motif, instead of YK in long pAgos, further supporting their separation into a distinct clade from long pAgos (Fig. 1, Fig. 2B, Fig. 3A and B). Though none of short pAgos was biochemically characterized to date, the conservation of key residues in the MID domain strongly suggests that these proteins bind 5’-phosphorylated guide nucleic acids similarly to most long pAgos and eAgos.

### Endonuclease activity of the PIWI domain

Structural and biochemical studies of several pAgos and eAgos demonstrated that the conserved tetrad of amino acid residues in the PIWI domain (DEDX, where X is D, H or K) is responsible for endonucleolitic cleavage of the target upon its recognition by the Ago-guide complex (Yuan et al. 2005; Wang et al. 2008b, 2009; Swarts et al. 2015a; Kaya et al. 2016). Similarly to previous reports (Makarova et al. 2009; Swarts et al. 2014b), we found that all pAgos that belong to the short clade lack a canonical DEDX catalytic tetrad in the PIWI domain (PIWI* variants) (Fig. 2A). Similar to short pAgos, all proteins in the long-B clade also lack the catalytic tetrad. In contrast, the majority of long-A pAgos (79%) have the canonical DEDX catalytic tetrad in the PIWI domain suggesting that they possess endonucleolytic activity. However, this branch also harbors pAgos with substitutions in the catalytic tetrad suggesting several independent events of the loss of endonuclease activity in this branch.

### Binding of the guide 3’-end in the PAZ domain

The amino acid sequences of the PAZ domains of the pAgo proteins are divergent, but their structural folds are similar. The full-length PAZ domain have two subdomains, each consisting of two nucleic-acid binding regions (Song et al. 2004; Yuan et al. 2005; Wang et al. 2008b; Liu et al. 2018), which are oriented to form a hydrophobic pocket that anchors the 3’-end of the guide strand in the binary pAgo-guide complex (the structure for TtAgo is shown in Fig. 4). The first subdomain consists of an oligosaccharide binding fold (OB-fold)-like structure with one or two helices on one side and includes nucleic acid binding regions R1 and R4. The second subdomain consists of an α-helix, a β-hairpin or loop structure, sometimes followed by another α-helix, and includes nucleic acid binding regions R2 and R3 (Fig. 4) (Song et al. 2004; Yuan et al. 2005; Wang et al. 2008b). In TtAgo, all four regions contribute to anchoring of the guide 3’-end in the PAZ pocket (Fig. 4) (Wang et al. 2008b). However, some pAgos, such as RsAgo and MpAgo, contain reduced variants of the PAZ domain that lack the second subdomain, including regions R2 and R3, and therefore do not possess the PAZ pocket (Kaya et al. 2016; Miyoshi et al. 2016; Liu et al. 2018). Nevertheless, structural analysis of the binary guide-MpAgo complex revealed that the guide 3’-end can still be bound in the incomplete PAZ domain, although with a different orientation relative to TtAgo (Fig. 4) (Kaya et al. 2016). These differences may possibly affect the kinetics of guide binding, the stability of binary guide pAgo complexes or their ability to recognize complementary nucleic acid targets (see Discussion).

**Figure 4.**
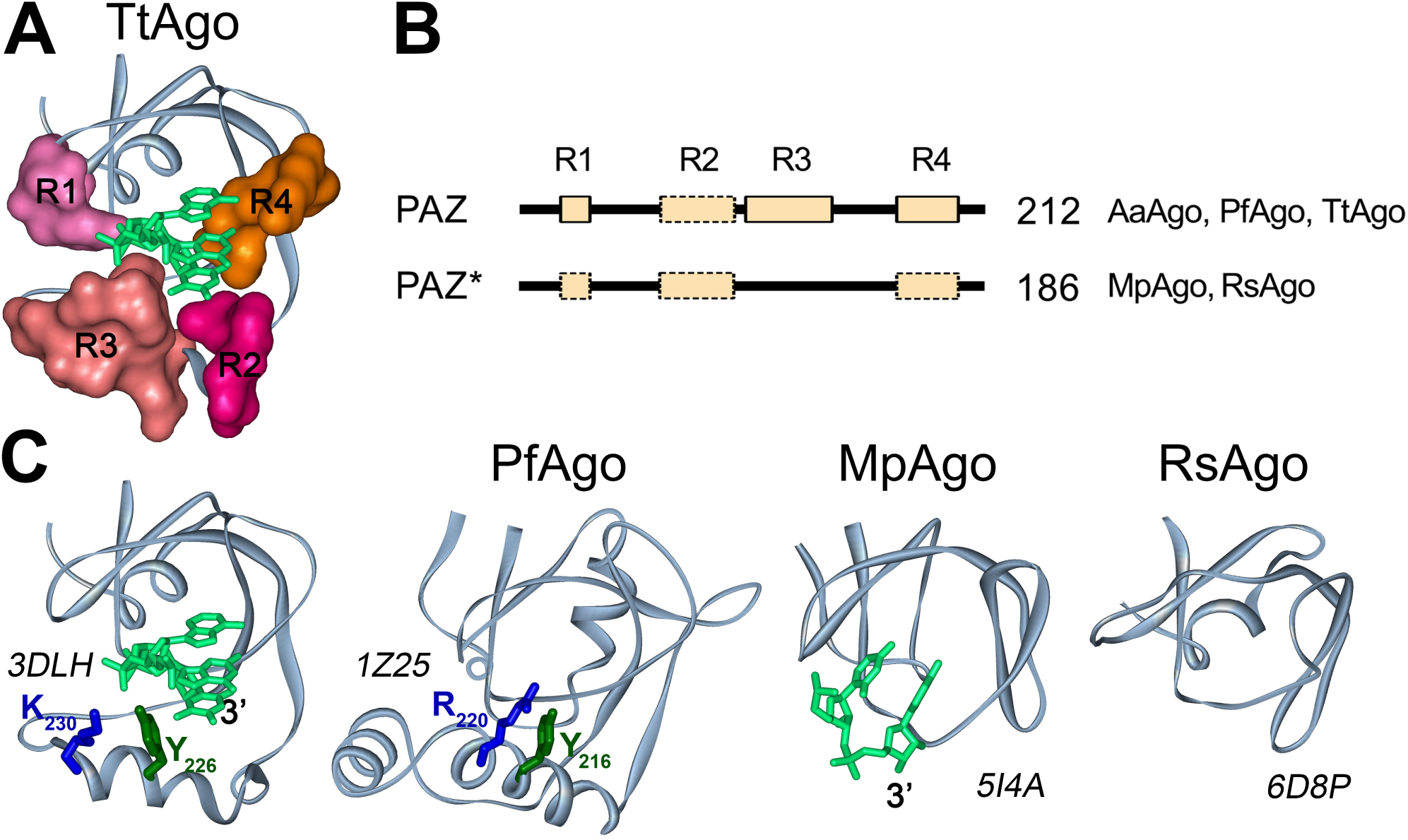
PAZ domains in the pAgo proteins. **(A)**. The structure of the 3’-guide binding pocket in TtAgo (PDB ID: 3DLH). Structural regions R1-R4 forming the pocket are shown in surface representation. **(B)**. A general scheme of the PAZ and PAZ* domain structures in long pAgos. All PAZ domains can have R1, R2, R3 and R4 regions. PAZ* is defined as variants of PAZ lacking full-length region R3. No regions R1, R2 and/or R4 could be detected in multiple alignments of some PAZ or PAZ* variants (shown by broken boxes), but their actual presence need to be tested due to their low conservation. The numbers show how many long pAgos have corresponding domain structure. The pAgos with resolved structures having PAZ and PAZ* domains are indicated. (**C)**. The three dimensional structures of PAZ (TtAgo, PfAgo) and PAZ* (MpAgo, RsAgo) variants in crystallized pAgos and their binary complexes with DNA (for TtAgo) or RNA (for MpAgo) guide molecules. Tyrosin and lysine/arginine residues from R3 in TtAgo and PfAgo, probably involved in guide binding, are indicated. PDB IDs are shown next to each structure.

We determined how often the PAZ domain without the second subdomain (referred to here as PAZ*) is present among long pAgos. As noted above, the structural elements of PAZ involved in nucleic acid binding are organized by the four poorly conserved regions, R1-R4 (Supplemental Fig. S3, as described in (Liu et al. 2018)), with the DNA-binding α-helix in the second subdomain formed by stretches of 7–15 amino acids from region R3 (illustrated for TtAgo and PfAgo in Fig. 4). Thus, we analyzed the presence of this region as a signature of the second PAZ subdomain in pAgos. Using multiple sequence alignments of the identified PAZ domains, we found that R3 is found in more than half of the PAZ domains in long pAgos, including most long-A pAgos (Fig. 1B, Fig. 4A), indicating their canonical two-lobe structure. The rest of PAZ* domains are smaller and do not have R3, suggesting that they have the one-subdomain organization without the nucleic-acid binding pocket, as previously observed for RsAgo and MpAgo. Interestingly, the PAZ* domain is found in almost all long-B pAgos (Fig. 1B), suggesting that it was present in the common ansestor of this group.

### Phylogenetic analysis of APAZ domain-containing proteins

Previous analyses identified the so-called APAZ domain that is present in some short pAgos as well as in genes encoded in the putative operons containing short pAgos (Makarova et al. 2009). The APAZ domain was proposed to be the functional analog (but not homolog) of the PAZ domain that is absent in short pAgos. To gain insight into the diversity and phylogeny of APAZ-containing proteins, we have iteratively searched the RefSeq protein database by DELTA-BLAST using the sequences of known APAZ domains as queries. Totally, we have found 5385 protein hits. However, the vast majorty (4753 or 88%) of the found proteins belonged to the ATP phoshorybosyltranspherase and phoshotranspherase EIIB families. We have performed the reverse DELTA-BLAST search using them as queries to test if that these protein families were indeed related to the other APAZ proteins, however, we did not find any APAZ-containing proteins within obtained hits. Moreover, the ATP phoshorybosyltranspherase proteins were found only among BLAST hits of the SIR2-APAZ protein from *Sphingomonas wittichii* RW1, while phoshotranspherase EIIB proteins were found only among BLAST hits of the TIR-APAZ protein from *Chlorobium phaeobacteroides* BS1. Thus, we concluded that ATP phoshorybosyltranspherase and phoshotranspherase EIIB families were revealed artificially and are not related to the APAZ proteins, and excluded them all from the further phylogenetic analysis. The remaining set of APAZ-domains included 632 proteins.

The multiple alignments of APAZ domains were used to build their phylogenetic tree that revealed that the APAZ proteins can be separated into four large groups designated Ia, Ib, IIa and IIb and a fifth group III of 18 proteins from several Archaean species that have remote similarity to groups I-II (Fig. 5). All five groups are present in putative operons with pAgos. We further analyzed the domain architectures of APAZ-containing proteins using InterProScan and the Pfam and Superfamily databases. The first group of APAZ proteins designated Ia includes 117 proteins that in addition to APAZ contain the SIR2-domain (the ‘SIR2-APAZ’ type). The majority of proteins in this group are short pAgos that also contain the MID-PIWI domains; however, 31 proteins from this group lack the MID-PIWI domains. Since these variants are scattered along different branches in this group, they might have independently lost their MID-PIWI part. In contrast, while a significant part of APAZ proteins from the four other groups are associated with pAgos, only one of them is pAgo itself. Most proteins that belong to group Ib (150) are related to group Ia and also contain SIR2 besides APAZ, but do not have MID and PIWI. There are also several proteins in both groups that lack SIR2 or any other domains (the ‘APAZ’ type; 4 and 6 proteins for Ia and Ib, correspondingly). Most proteins of group IIa (122) are characterized by the presence of the TIR domain on their N-termini (‘the TIR-APAZ’ type); one of them is short pAgo (see also Fig. 2); 16 proteins in this group lack TIR domains. Group IIb, which encompasses 225 proteins, is diverse: 112 proteins contain uncharacterized domain DUF4365 (‘DUF4365-APAZ’), 13 have the SIR2 domain, while 100 do not have any extra domains (‘APAZ’). Our analysis of the DUF4365 domain have shown that it corresponds to the previously identified novel RecB-like domain of the Mrr subfamily of PD-(D/E)XK nucleases (see next section) (Makarova et al. 2009). Finally, the proteins from group III, similarly to a major part of proteins from group IIb, do not have any other domains except APAZ (the ‘APAZ’ type).

**Figure 5.**
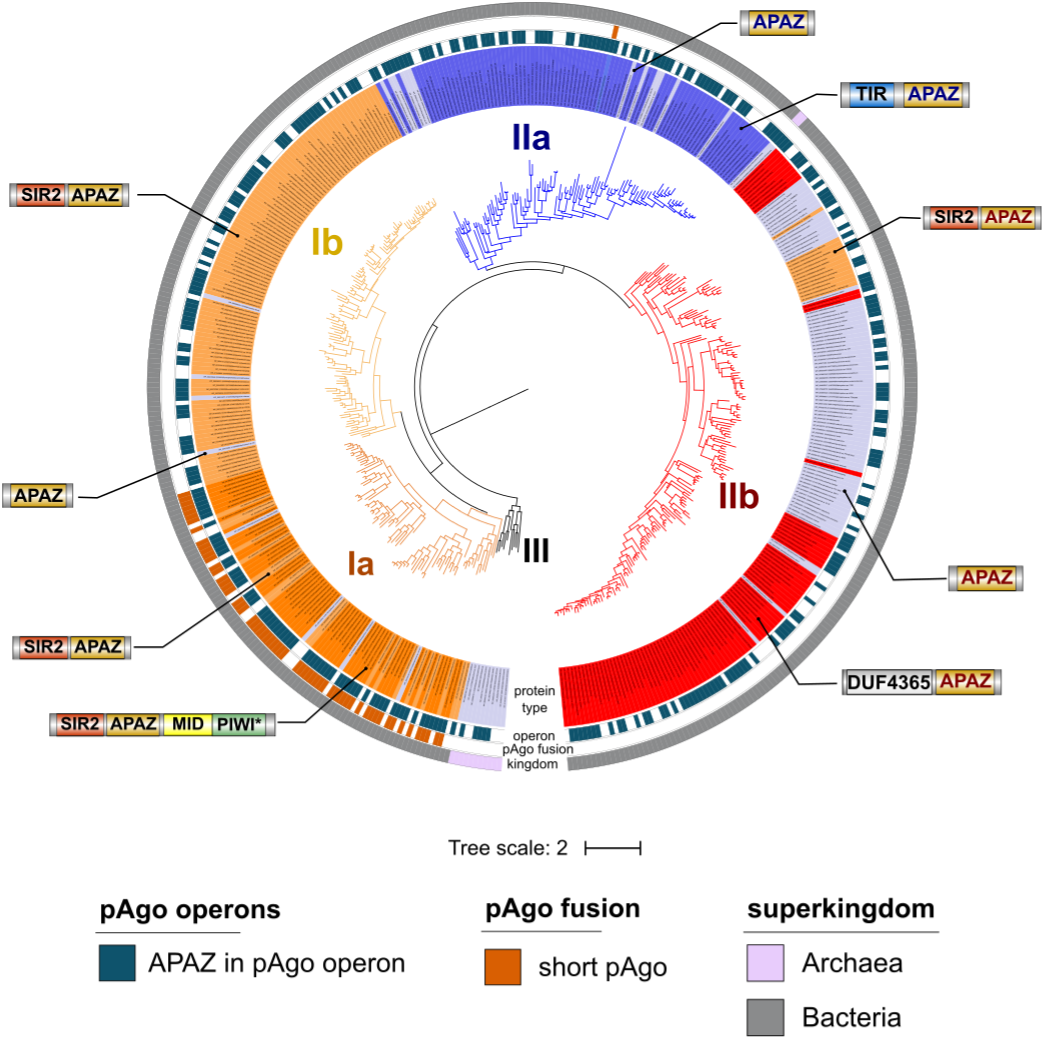
Phylogenetic analysis of APAZ domains. The circular phylogenetic tree of the five groups of APAZ domains. The phylogenetic tree was annotated as follows, from the inner to the outer circles: in the ‘protein type’ circle the APAZ-containing proteins are colored according to their phylogenetic groups and domain compositions, where each type of protein structure is exemplified by the callouts with corresponding domain schemes; isolated APAZ domains in all groups are shown in light blue; the ‘operon’ circle indicates whether the APAZ-containing protein is found in a pAgo coding operon; the ‘pAgo fusion’ circle shows APAZ domains which are fused with short pAgo proteins; the ‘kingdom’ circle indicates the superkingdom to which the corresponding APAZ-containing protein belongs to.

APAZ-containing proteins are found almost exclusively in Eubacterial species. The analysis revealed tight reciprocal association between APAZ-containing proteins and short pAgos. Indeed, although only one branch of short pAgos contain the APAZ domain, the majority of other short pAgos that do not have APAZ themselves encode APAZ-containing genes in their operons (Fig. 1B, Fig. 2). Reciprocally, 397 or 63% of all genes with the APAZ domain are positioned close (within 10 genes on the same genomic strand) to short pAgos, likely in the same operons (Fig. 5) (see next section). Thus, APAZ proteins are likely an integral component of functional pathways mediated by short pAgos.

### Functional classification of proteins enriched in the genomic context of pAgos

Analysis of the functions of proteins encoded in the same operons with pAgos might shed light on the molecular pathways involving pAgos. We therefore analyzed the genomic context of pAgos encoded in the 1385 genomes of 1248 strains and explored the proteins encoded in their proximity. We defined a window centered on pAgo encompassing 20 genes (*i.e*. ten genes upstream and ten genes downstream of pAgo) in each genome. The 16274 proteins encoded in these windows were clustered into orthogroups based on the sequence similarities that resulted in 1892 orthogroups that had at least 2 proteins (see Methods). The orthogroups contained 12668 proteins and the largest one comprised 298 related proteins (Supplemental Table S2). We further defined pAgo operons that included only genes that are continuously located on the same genomic strand as pAgo that resulted in a set of operons with mean size of 6.4 genes. We separately considered operons of five different groups of pAgos (Fig. 6): (1) short pAgos that are not fused to the APAZ domain; (2) short pAgos that contain the SIR2-APAZ domains; (3) all long-A pAgos except the MID-OH variants; (4) long-B pAgos; and (5) long-A pAgos with the MID-OH domain. Proteins that belonged to 47 orthogroups were present in at least 5% of operons of any of the five pAgo groups and were further analyzed in detail. In Fig. 6, the orthogroups are sorted in descending order of the number of proteins in each orthogroup.

**Figure 6.**
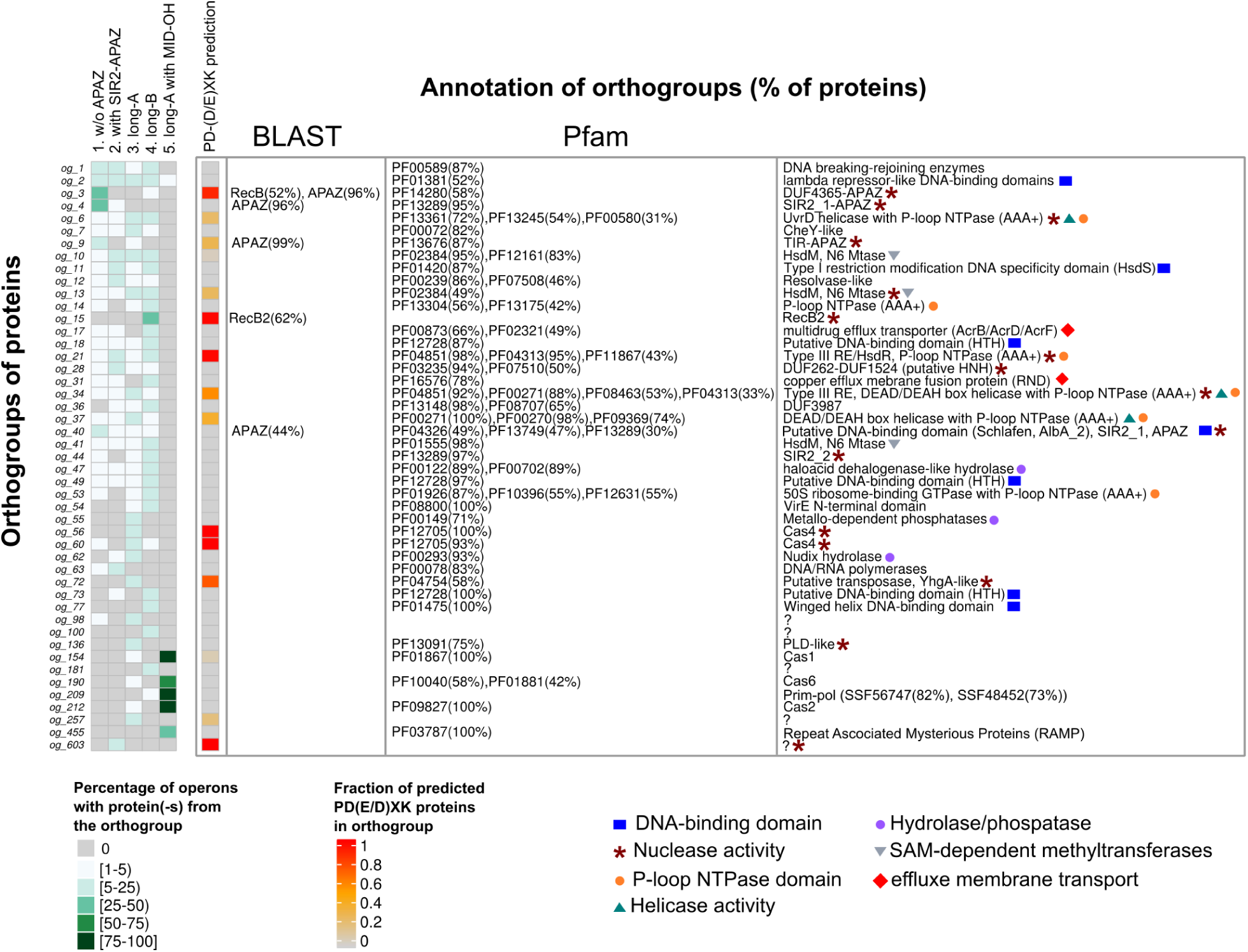
Functional annotations of proteins encoded in pAgo operons. The heatmap on the left shows the percentage of operons for each pAgo group (columns) that have at least one protein from the particular orthogroup (rows). The orthogroups are ordered according to the number of proteins in each group, from the largest to the smallest. The PD-(D/E)XK column shows the fraction of proteins from each orthogroup that are predicted to be nucleases from the PD-(D/E)XK superfamily. The annotation of proteins was performed by BLAST analysis for identification of APAZ, RecB-like, RecB2 domains and by InterProScan with using Pfam and Superfamily databases. Only the Pfam annotation of domains is shown (with the exception of og_209 for which the Superfamily annotation is shown). The percentage in the brackets after the domain name shows the fraction of proteins in the corresponding orthogroup having this type of domain. Only Pfam or Superfamily domains found in at least 30% of orthogroup proteins are shown. The descriptions of domain functions are shown on the right of the panel. Proteins from several orthogroups do not have any known domains and are marked by “?”. After the manual curation of domain functions, several most common functional features were selected, which are shown as colored symbols and described below the table.

We defined the domain structure of proteins in each orthogroup using InterProScan with Pfam, Superfamily and CDD databases containing known domain annotations. For identification of the APAZ, RecB-like and RecB2 domains, previously described by Makarova et al. and absent from these databases, we performed PSI-BLAST search using previously identified proteins as queries. In addition, we proposed that there may be other pAgo-associated proteins belonging to the PD-(D/E)XK superfamily of nucleases, besides previously identified Mrr (RecB-like and RecB2) and Cas4 proteins (Makarova et al. 2009; Swarts et al. 2014b). To check this, we have searched for the characteristic PD-(D/E)XK sequence motif in the orthogroup proteins. The identified domains found in each orthogroup of proteins and the putative functional types of pAgo-associated proteins are shown on Fig. 6.

The proteins with APAZ domains are present in the orthogroups og_3, og_4, og_9 and og_40. The proteins from these orthogroups also contain other types of domains, including DUF4365 (og_3), SIR2_1 subtype of SIR2 (og_4), TIR (og_9), and SIR2_1 fused with Schlafen domain (og_40). Most of revealed APAZ-containing proteins are located in group 1 operons of short pAgos that do not have their own APAZ domain. Totally, 91% of group 1 operons contain proteins with APAZ. In contrast, only 6% of group 2 operons of short pAgo fused with APAZ and SIR2 encode APAZ containing proteins. Taken together, 93% of short pAgos are associated with APAZ domain as a part of the same protein or present in the same operon with short pAgos. In contrast, long pAgos only rarely co-encoded with APAZ proteins (2%, 3% and 0% of operons from groups 3, 4 and 5 respectively).

We found several different types of nucleases and DNA-binding proteins abundant in the pAgo operons (Fig. 6). Totally, 17 out of 47 orthogroups (36%) encode proteins with putative nuclease activity. These proteins are found in 68% of pAgo operons. 92% of operons of short pAgo that themselves do not have the APAZ and SIR2 domains (group 1 of pAgos) encode different types of nucleases. In contrast, only 32% of operons of short pAgo fused with the SIR2 and APAZ domains (group 2) have nucleases. For group 3 of long-A pAgos, 45% pAgos carrying the catalytic tetrad have nucleases in their operons, while this number is increased to 53% for operons of long-A lacking the DEDX tetrad. This number is further increased to 67% for group 4 of long-B pAgos, which are all catalytically inactive. The nucleases found in the pAgo operons are quite diverse and include proteins with SIR2 (og_4, og_40 and og_44), TIR (og_9) and RecB2 (og_15) domains, the DUF4365 domain that belongs to RecB-like nucleases (og_3), Cas4 proteins (og_56 and og_60) and phospholipase D proteins with putative nuclease activity (og_136). A set of pAgo operons also contain proteins with putative nucleases of the HNH type with DUF252 and DUF1254 domains (og_28).

Two pAgos from *Marinitoga piezophila* and *Thermotoga profunda*, which are representatives of the small group of the long-A pAgos having the MID-OH domain with specificity to 5’-OH ends of guide molecules (Fig. 1B, Fig. 2B, Fig. 3A), were previously reported to be encoded in the CRISPR loci between the Cas1 and Cas2 genes (Kaya et al. 2016). We found that all pAgos contaning the MID-OH domain from 9 prokaryotic strains (group 5 of pAgos) are encoded within CRISPR loci with Cas1 (og_154) and Cas2 (og_209) genes nearby (Fig. 6). Also, there are other proteins from the CRISPR locus belonging to subtype III-B observed in the operons of MpAgo-like pAgos: Cas6 (og_190), primase (og_209) and Cmr (Cas module RAMP) proteins (og_455).

In addition, the proteins in several other orthogroups have motifs characteristic for the PD-(D/ E)XK nucleases (Fig. 6, the PD-(D/E)XK column). These proteins include the UvrD helicase and P-loop NTPase (AAA+ subfamily) domains (og_6), SAM-dependent methyltranspherase domain (og_13), restriction endonuclease of Type III (og_21, og_34), DEAD-box helicase and P-loop NTPase (AAA+ subfamily) (og_34 and og_37), YhgA-type putative transposases (og_72) and proteins without known domains (og_257 and og_603). Notably, the proteins in three of the orthogroups that contain PD-(D/E)XK motifs have predicted helicase activity as they possess domains related to the UvrD helicase (og_6) and DEAD-box helicase (og_34 and og_37).

Proteins with DNA-binding domains are abundant in operons of short pAgos fused to the APAZ and SIR2 domains (33%) and of long-B pAgos (33%). In addition, proteins with DNA-binding domains are present in 19% and 13% operons of long-A pAgos and short pAgos without APAZ-SIR2 domains, respectively. The DNA-binding domains in pAgo-associated proteins include the lambda repressor-like DNA-binding domain (og_2), helix-turn-helix DNA-binding domains (og_18, og_49, og_73 and og_79), the Schlafen domain with an AlbA_2 subdomain (og_37) and DNA-binding components of the Type I restriction-modification system (HsdS) (og_11). Interestingly, the lambda repressor-like DNA-binding domain (PF01381) from highly represented orthogroup og_2 and the Schlafen DNA-binding domain AlbA_2 (PF04326) from og_37 were found to be fused with several long-B pAgos (Fig. 2B).

Overall, analysis of proteins in pAgo operons strongly indicate the involvement of nuclease, helicase and DNA-binding activities in cellular pathways dependent on pAgos (see Discussion). Beyond nucleases and DNA-binding proteins, 10% of all long pAgo operons encode hydrolases and phosphatases (og_47, og_62, og_55). Finally, 14% of long-B pAgo operons encode multidrug efflux transporters of the resistance-nodulation-cell division (RND) superfamily (og_14 and og_31), but their possible roles in the functional pathways involving pAgos remain unknown.

### The genomes encoding pAgos do not show decreased transposon copy number

Several eukaryotic Ago proteins were shown to prevent propagation of transposons and viruses in the genomes (reviewed in (Juliano et al. 2011; Hutvagner and Simard 2008)). Similarly, a few studied pAgos were proposed to protect prokaryotic cells from foreign DNA, including plasmids, transposons and phages (Olovnikov et al. 2013; Swarts et al. 2015a; Zander et al. 2017; Swarts et al. 2015a). In particular, pAgos may be involved in the protection of the prokaryotic genome from the invasion of mobile genetic elements or suppress the activity of genomic copies of such elements. If pAgos are able to reduce the expansion of transposons in their host genomes then the number of transposon insertions should be lower in strains that encode pAgo. To test this hypothesis, we counted the number of transposon insertions in the genomes of strains encoding or do not encoding pAgos from different groups (short pAgos, short pAgos fused to the SIR2-APAZ domains, long-A, long-B pAgos, and long-A pAgos with the MID-OH domain). We found that the genomes of strains encoding pAgos do not have lower copy number of transposons in comparison to the strains without pAgo genes (Fig. 7). Moreover, we observed a moderate but significant increase in the mean number of transposon copies per genome in pAgo-contaning strains for both short and long classes of pAgos. For the MID-OH pAgos, the number of transposon copies in corresponding genomes was smaller than in pAgo-minus strains, but this difference was statistically insignificant due to the small number of pAgos in this group. Overall, our results indicate that pAgos present in the genomes may not be able to stop transposon insertions, however, it does not exclude a possibility that pAgos can repress transposon expression after their integration.

**Figure 7.**
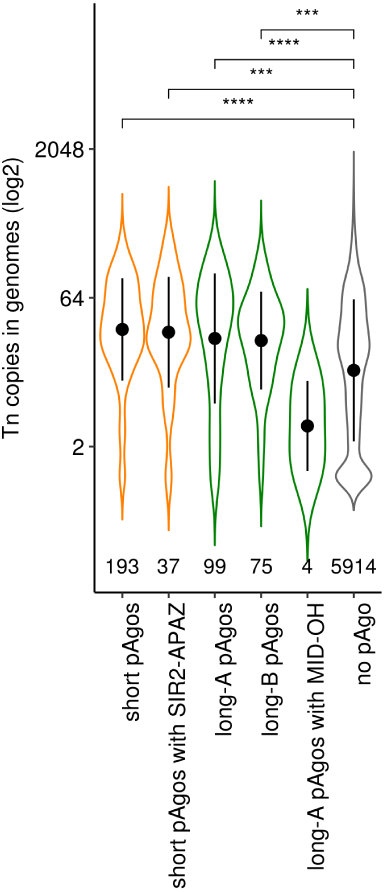
The number of transposon copies in genomes with and without pAgo genes. The violin plot shows the distribution of copy numbers of transposons in the genomes of prokaryotic strains encoding or do not encoding pAgos. The black dots with whiskers indicate the means and standard deviations. The numbers of analyzed genomes for each group of pAgos are shown below each violin. The p-values shown above the plot were calculated with the Wilcoxon ranked test: **** p-value ≤ 1×10^−4^, *** p-value ≤ 1×10^−3^; only p-values ≤ 0.05 are shown.

## Discussion

Eukaryotic RNAi systems have evolved from several diverse prokaryotic components including RNaseIII-like enzymes, RNA-dependent RNA polymerases, helicases and pAgos, which gave rise to various types of eAgos and RNAi pathways (Koonin 2017). While being highly variable and involved in different interference pathways, all eAgos share the same *modus operandi*: they always bind small RNA guides to locate and inactivate complementary RNA targets (or attract other components to modify corresponding coding DNA loci). Perhaps not surprising, pAgos are much more diverse, with significant sequence and domain variations (Makarova et al. 2009; Swarts et al. 2014b). Moreover, a few published functional studies of pAgos demonstrated that, while retaining the same basic principle of guided nucleic acid recognition, they may significantly differ in their modes of action in comparison with eukaryotic counterparts (Olovnikov et al. 2013; Swarts et al. 2014a; Zander et al. 2017; Swarts et al. 2015b, 2015a). We have now revealed even a larger diversity of pAgos that allows to suggest their updated classification, propose possible evolutionary pathways and cellular functions for this conserved family of nucleic-acid-targeting proteins.

### The Diversity of pAgos

Our results revealed that pAgos are widespread in Archaean and Eubacterial species. The availability of new sequenced genomes allowed identification of an almost 3-fold larger set of pAgos compared to previous analysis. Analysis of this expanded set demonstrated that pAgos are highly diverse and can be separated into three large clades: short, long-A and long-B that diverged early in evolution (Fig. 1B; Supplemental Fig. S1). While this division is based solely on sequence alignments of MID-PIWI domains, these groups also reveal characteristic variations in specific structural elements involved in nucleic acid binding and processing. Thus, the members of the long-A clade mostly contain both catalytically active PIWI domain and PAZ domain of normal size. In contrast, long-B pAgos contain inactive PIWI* domain and also usually shortened PAZ* domain, although other combinations were also observed for both clades (*e.g*. PAZ-PIWI* for both long-A and long-B pAgos, Fig. 1 and 2). Furthermore, proteins from the long-A clade contain several variants of the 5’-P-guide binding motif (see below) while long-B pAgos almost always contain the canonical YK motif. Similarly, short pAgos always contain substitutions in the active site (PIWI*) and a subset of canonical MID motifs (HK/RK).

The exact relationship between the three clades is ambiguous with two possible alternative scenarios (Swarts et al. 2014b). In one scenario, short and long pAgos diversified first, followed by further split of long pAgos into two clades. Alternatively, catalytically inactive long-B pAgos and short pAgos may have originated from a common ancestor containing substitutions in the active site. The phylogenetic analysis of the MID-PIWI domains has shown that the evolutionary distance between combined short pAgo and long-B pAgo clades on the one side and the long-A clade on the other is slightly larger than between short and combined long-A and long-B clades (Supplemental Fig. S1). This argue in favor of the second scenario of the evolutionary origin of the three clades of pAgos.

### Binding of nucleic acids by pAgos

Tight association with small nucleic acid guides seems to be a defining feature of pAgos and eAgos alike (reviewed in (Swarts et al. 2014b; Lisitskaya et al. 2018)). Analysis of the nucleic acid binding pockets of the MID domain revealed that the majority of pAgos that belong to all three clades (short, long-A and long-B) likely bind small nucleic acids guides with monophosophorylated 5’-ends. At the same time, we discovered significant variations in the conserved 5’-guide binding motifs in the long-A pAgos, including substitutions of key residues involved in the 5’-phosphate interactions (*i.e*., YK, YY, KR, YR, MID*, MID-OH variants). While most these variants likely interact with 5’-P-guides, some proteins might bind guides with an alternative 5’-end structure, as already shown for the MID-OH pAgos, or not use nucleic-acid guides at all. Interestingly, almost all members of the short pAgo clade have MID variants that are close to the consensus (HK/RK), suggesting that the canonical MID pocket structure is essential for their function in the absence of the PAZ domain. Future biochemical comparisons of the long and short pAgos with various noncanonical MID motifs should find out if the observed differences in the 5’-end pockets may influence the specificity, affinity and kinetics of their interactions with guide and target molecules.

While all known eAgos use small RNA guides to recognize RNA targets, the majority of proteins in a relatively small sample of pAgos characterized so far prefer to bind small DNA as a guide, with the exception of RsAgo(Olovnikov et al. 2013) and MpAgo (Kaya et al. 2016) that associate with small RNAs. Furthermore, all studied pAgos preferably interact with DNA targets. Few amino acid residues involved in the binding of ribose- and deoxyribose-backbone of the guide and target molecules were identified in the structures of experimentally characterized pAgos (Wang et al. 2008a; Sheng et al. 2014; Kaya et al. 2016; Miyoshi et al. 2016; Liu et al. 2018), however, these residues are not well conserved. Therefore, sequence analysis is not able to predict preference for DNA *vs* RNA binding, and this information has to be obtained experimentally.

The PAZ domain of most studied eAgo and pAgo proteins is composed of two subdomains that together form the 3’-end guide binding pocket (Song et al. 2004; Yuan et al. 2005; Wang et al. 2008b; Liu et al. 2018). However, our results revealed different organization of the PAZ domain among long pAgos indicating that a single conserved structure of PAZ may not be required for guide binding. The structure is not conserved even among few experimentally characterized pAgos. While AaAgo and TtAgo contain both subdomains (Yuan et al. 2005; Wang et al. 2008b), the PAZ* domains of MpAgo and RsAgo lack the second subdomain, and therefore do not possess the PAZ pocket (Kaya et al. 2016; Miyoshi et al. 2016; Liu et al. 2018). While no three dimensional structure of the binary complex is available for RsAgo, the guide 3’-end still interacts with PAZ in the binary complex of MpAgo but is differently oriented (Fig. 4) (Kaya et al. 2016). Formation of the extended guide-target base-pairing during target recognition is accompanied by guide release form PAZ (Wang et al. 2009; Sheng et al. 2014; Zander et al. 2014; Kaya et al. 2016; Doxzen and Doudna 2017). Thus, structural characteristics of PAZ may have an important role in the recognition and binding of nucleic acid targets by pAgo proteins, for example by influencing the kinetics of target recognition and cleavage (Wang et al. 2009; Jung et al. 2013; Sheng et al. 2014; Zander et al. 2014; Kaya et al. 2016; Doxzen and Doudna 2017). Interestingly, the reduced PAZ* domain is characteristic for the long-B clade of catalytically inactive pAgos, suggesting that the stepwise guide-target annealing controlled by full-length PAZ may be more important in the active long-A pAgos, *e.g*. for prevention of premature nonspecific target cleavage.

All short pAgos and a small, but significant fraction of long pAgos lack the PAZ domain entirely raising the question of how nucleic acid guides can be bound by these proteins. Structural and functional analysis of the truncated long-A AfAgo lacking the PAZ domain demonstrated that it can nevertheless bind and stabilize short guide-target duplexes (Parker et al. 2004, 2005, 2009). Furthermore, since the guide 3’-end is released from the PAZ domain upon target recognition, PAZ may be important for protection of the guide from cellular nucleases until the target is found – and the protection may be less important for short pAgos or truncated long pAgos that use DNA guides. However, it remains to be established whether any accessory proteins may facilitate guide and target binding by such pAgos. The finding of association between short pAgos and the APAZ domain (either in the same protein or encoded by a separate gene in the same operon) led to the suggestion that APAZ might play the role of PAZ in short pAgos (Makarova et al. 2009). Recently, it was proposed that APAZ may be instead homologous to the N-domain of full-length Agos (Willkomm et al. 2018). However, both proposals await experimental validation.

### Possible functional activities of pAgo-associated proteins

In contrast to their ability to bind nucleic acids, the endonuclease activity towards the target is not a universal feature of pAgos: the proteins that are predicted to have the catalytic tetrad in the PIWI domain are found in only the long-A clade of pAgos (~18% of all pAgos). However, the lack of catalytic activity in PIWI* does not rule out the possibility that such pAgos are involved in processing of target nucleic acids. Indeed, our analysis confirmed and expanded previous observations on the association of catalytically inactive pAgo with proteins that might have nuclease activity (Makarova et al. 2009; Swarts et al. 2014b). Markedly, the number of putative nuclease genes in operons containing catalytically inactive pAgo variants is significantly higher than for the active pAgos.

The proteins in one branch of short pAgos have the SIR2 domain with potential nuclease function, while many more pAgos that belong to the short and long-B clades of catalytically inactive pAgos harbor in their operons separate proteins with predicted nuclease activities. The absence of the catalytic tetrad in the PIWI* domain in such pAgos and their tight association with proteins containing SIR2, TIR2, DUF4365 (novel RecB-like nuclease of the Mrr subfamily), RecB2, Cas4, PLD, HNH nuclease domains suggest that these domains might replace the intrinsic endonuclease activity of the Ago protein. Although the APAZ domain is not related to any nuclease, its strong association with short pAgos and the proteins with nuclease domains suggests that APAZ might also be involved in the nuclease activity. Thus, the functions of target recognition and its further processing seem to be separated, with the first step performed by pAgos and the second by nucleases encoded in the pAgo operons. Notably, these operons also include predicted helicases and DNA-binding proteins, suggesting their involvement in DNA processing. Future analysis of the functional activities of these proteins will be required to uncover the molecular details of pAgo action.

### Possible functions of pAgos in prokaryotic cells

eAgos and their small RNA partners play diverse functions, from the repression of selfish genomic elements such as transposons to the regulation of host gene expression. Furthermore, eAgos can direct changes of chromatin marks and even DNA elimination (Hutvagner and Simard 2008; Juliano et al. 2011). Compared to eAgos, experimental data about pAgo functions are very scarce, as activities of only a handful of proteins were characterized *in vitro* and even smaller number of proteins (TtAgo and RsAgo) were studied in their host cells *in vivo*. Available data suggest that pAgo might participate in protection of prokaryotic genomes against foreign elements such as plasmids, transposons and phages (Olovnikov et al. 2013; Swarts et al. 2014a). However, the only direct results obtained so far was the ability of TtAgo and RsAgo (and also PfAgo and MjAgo in the heterologous *E. coli* system) to decrease the amount of plasmid DNA in the cell (Olovnikov et al. 2013; Swarts et al. 2015a, 2014a; Zander et al. 2017). RsAgo was also demonstrated to decrease plasmid transcription and preferentially associate with genomic DNA sequences corresponding to transposons and IS elements (Olovnikov et al. 2013). While the molecular mechanisms of this specific targeting remain to be established, we infer that other types of pAgos discovered in our analysis may also be involved in host defense against certain types of genetic elements.

Intriguingly, we observed a positive, not negative, correlation between the presence of various types of pAgos and the number of transposon copies in sequenced genomes. This observation suggests several possibilities. First, transposons may not be the main targets of all pAgos, most of which may instead mainly target phages and plasmids. Second, the presence of pAgos in genomes with increased transposon numbers might be beneficial for these strains because pAgos suppress further detrimental transpositions. Third, it can be speculated that pAgos might perform a plethora of other functions in prokaryotic cells, including genetic regulation and DNA repair. Indeed, association of pAgos with multiple DNA-binding and DNA-processing proteins suggests their active involvement in DNA metabolism in prokaryotic cells.

As reported previously (Makarova et al. 2009; Swarts et al. 2014b) and in this work, the pattern of distribution of pAgos among Eubacterial and Archaeal species does not correspond to the phylogeny of host species suggesting their spread by horizontal gene transfer. Other genes spread by HGT include various adaptation and resistance systems such as CRISPR, restriction/modification, drug resistance, toxin/antitoxin modules that have common features of cell protection in the highly diverse and developing environments. While it is attractive to suggest that most pAgos use associated nucleic acid guides for sequence specific recognition of foreign DNA targets in order to repress them, other scenarios of their action are as well possible. For example, pAgos together with their protein partners might constitute suicidal system that triggers processing of host DNA unless its activity is suppressed. Our study opens new possibilities for detailed analysis of various groups of pAgos and associated factors, which can reveal their new cellular functions and activities and possibly adopt pAgos as an efficient tool for genomic manipulations.

## Methods

### Prokaryotic Protein and Genome Databases

The set of proteins were downloaded from the NCBI FTP site in January 2018. Altogether the database included more than 116 million RefSeq proteins from 4,364 taxons with completely sequenced and assembled genomes (“Complete Genome” and “Chromosome” statuses) and 15,505 taxons with unassembled genomic sequences (“Scaffold” and “Contig” statuses) of Archaea and Eubacteria. The genomic sequences and genome annotations of prokaryotes were fetched from the NCBI FTP site in January 2018.

### Identification of pAgo and APAZ-containing proteins and analysis of their domain structure

The identification of homologs of already known pAgo- and APAZ domain-containing proteins was carried out using the PSI-BLAST and DELTA-BLAST program from the NCBI-BLAST+ package, v.2.6.0 (`-num_descriptions 1000 -num_alignments 1000 -evalue 10E-5 - num_iterations 5`) (Altschul et al. 1997). The search was performed with five iterations, sufficient for the full convergence of the results. For the search, sequences of only the PIWI-MID domains of already known pAgos (Swarts et al. 2014b) or APAZ domains (Makarova et al. 2009) were used as queries. The domain architecture of the found proteins was analyzed by the CDD-batch program (https://www.ncbi.nlm.nih.gov/Structure/bwrpsb/bwrpsb.cgi) for the Pfam and CDD databases as well as InterProScan (v.5.28-67) for the Pfam and Superfamily databases. The proteins were aligned by the MAFFT program, v.7.3 (`-ep 0 --genafpair --maxiterate 1000`) (Katoh and Standley 2013); miltiple alignments were manually curated and used for the extraction of domain borders and features.

### Phylogenetic Analysis

To construct a non-redundant, representative sequence set for the phylogenetic analysis, sequences of the PIWI-MID and APAZ domains were clustered using the UCLUST 4.2 program (Edgar 2010) with the sequence identity threshold of 90%. The longest sequence was selected to represent each cluster. The multiple alignment of PIWI-MID domains was carried out by the MAFFT program, v. 7.3 (-ep 0 --genafpair --maxiterate 1000) (Katoh and Standley 2013). Positions including greater than >=0.5 gaps were removed from the alignment by trimAl, v.1.4 (Capella-Gutiérrez et al. 2009). Phylogenetic analysis was performed using the FastTree program (Price et al. 2010) with default parameters, with the WAG evolutionary model and the discrete gamma model with 20 rate categories. The tree structure was validated with bootstrap analysis (n=100).

### Analysis of the composition of pAgo operons

The protein-coding sequences of 16264 genes neighboring all pAgo genes (ten upstream and ten downstream genes for each pAgo) were clustered based on sequence homology. The pairwise similarity was calculated by using the profile-profile metric with HHblits and HHsearch from the HH-suite (Soding 2005; Remmert et al. 2012), as suggested in (Bernardes et al. 2015). In contrast to the more common sequence-sequence comparison method with using all-against-all BLAST analysis, the comparison of HMM profiles represents a more sensitive strategy for detecting distant evolutionary relationships among diverged proteins. For this, a profile for each sequence was constructed with HHblits (`-n 1 -E 1E-3`) and *uniclast30_2017_10* HMM database (http://wwwuser.gwdg.de/~compbiol/uniclust/2017_10). Then, all-against-all profile–profile search was conducted using HHsearch (`-b 1 -b 1000 -z 1 -Z 1000 -E 1E-5`), which provided e-values similar to what BLAST does. Clustering of genes into orthogroups was done with the MCL algorithm implemented in *clusterMaker2* plug-in of Cytoscape 3.6.1 (inflation factor = 2).

The annotation of obtained 1892 protein orthogroups was perfomed with InterProScan (v.5.28-67) for the Pfam and Superfamily databases. Also we have performed the PSI-BLAST search of APAZ, RecB-like and RecB2 domains in the orthogroup proteins. The prediction of nucleases belonging to the PD-(D/E)XK superfamily was carried out with SVM-based approach using the standalone version of ‘PDEXK recognition’ program kindly provided by Dr. Mindaugas Margelevičius (Laganeckas et al. 2011). For each pAgo operon we then determined the orthogroups to which operon genes belonged to. If a pAgo was encoded in several genomes, only the genome with the largest operon was selected for final counting. Only orthogroups that were present in at least 5% of operons for each pAgo type (minimum three operons of any type of pAgo operon) were kept.

### Counting the transposon copy number in genomes

The transposon insertions in genomes were counted using genomic annotations of prokaryotic strains fetched from NCBI. Only genomes with statuses ‘Complete Genome’ and ‘Chromosome’ were taken into account. For each genome we counted the number of occurrences of key words ‘insertion sequence’ or ‘transposase’ in the genomic annotation file. If there were several completed genomes for a given strain then the median number of transposon insertions was calculated. If a pAgo gene was found in different strains of the same species (having the same taxon_id in the NCBI annotation), then the median number of transposon insertions among all strains was taken.

## Acknowledgments

The authors thank Mindaugas Margelevičius for the help with the prediction of PD-(D/E)XK motifs in proteins. This work was supported by the Grant of the Ministry of Education and Science of Russian Federation 14.W03.31.0007.

## Supplemental Information

**Supplemental figure S1.** The correlation between the overall number of analyzed genera with sequenced genomes in prokaryotic classes with the number of genera encoding pAgos. The size of points reflects the number of found pAgo genes in each class. The regression line is shown in blue; the grey zone around the regression line is the 95% confidence interval. The Pearson correlation coefficient and p-value are shown.

**Supplemental figure S2.** The unrooted phylogenetic tree of the PIWI-MID domains. The branches were collapsed into triangles, the size of which is proportional to the number of collapsed nodes. The numbers under the branches are the bootstrap support values (shown as fractions, n=100).

**Supplemental figure S3.** A fragment of the multiple alignment for selected PAZ and PAZ* domains. Regions R1-R4 that correspond to the structural elements involved in 3’-end guide binding are highlighted by yellow rectangles.

**Supplemental table S1:** The list of identified pAgo proteins.

**Supplemental table S2.** The summary of protein orthogroups. For each orthogroup the included in proteins and total number of proteins are shown.

**Supplemental File 1**. The alignments of proteins and domains described in this study (ZIP archive). Content: 1) The alignment of MID* domains that do not have one or more conserved amino acid resuides responsible for the binding of 5’-P or 5’-OH ends of the guide molecule. Only a fragment of the alignment of MID* corresponding to the 5’-end binding motif in MID is present. 2) The alignment of PAZ and PAZ* domains of long pAgos. 3) The alignments of five groups of APAZ domains that were used for the phylogenetic analysis. Positions including greater than >=0.5 gaps were removed from the alignments.

## References

Altschul SF, Madden TL, Schaffer AA, Zhang J, Zhang Z, Miller W, Lipman DJ. 1997. Gapped BLAST and PSI-BLAST: a new generation of protein database search programs. Nucleic Acids Res 25: 3389–3402.

Behm-Ansmant I. 2006. mRNA degradation by miRNAs and GW182 requires both CCR4:NOT deadenylase and DCP1:DCP2 decapping complexes. Genes Dev 20: 1885–1898.

Bernardes JS, Vieira FR, Costa LM, Zaverucha G. 2015. Evaluation and improvements of clustering algorithms for detecting remote homologous protein families. BMC Bioinformatics 16: 34.

Bujnicki JM, Rychlewski L. 2001. Identification of a PD-(D/E)XK-like domain with a novel configuration of the endonuclease active site in the methyl-directed restriction enzyme Mrr and its homologs. Gene 267: 183–191.

Capella-Gutiérrez S, Silla-Martínez JM, Gabaldón T. 2009. trimAl: a tool for automated alignment trimming in large-scale phylogenetic analyses. Bioinformatics 25: 1972–1973.

Djuranovic S, Nahvi A, Green R. 2012. miRNA-Mediated Gene Silencing by Translational Repression Followed by mRNA Deadenylation and Decay. Science 336: 237–240.

Doxzen KW, Doudna JA. 2017. DNA recognition by an RNA-guided bacterial Argonaute. PLoS One 12: e0177097–e0177097.

Edgar RC. 2010. Search and clustering orders of magnitude faster than BLAST. Bioinformatics 26: 2460–2461.

Elkayam E, Kuhn CD, Tocilj A, Haase AD, Greene EM, Hannon GJ, Joshua-Tor L. 2012. The structure of human argonaute-2 in complex with miR-20a. Cell 150: 100–110.

Faehnle CR, Elkayam E, Haase AD, Hannon GJ, Joshua-Tor L. 2013. The making of a slicer: activation of human Argonaute-1. Cell Rep 3: 1901–1909.

Hegge JW, Swarts DC, van der Oost J. 2017. Prokaryotic Argonaute proteins: novel genome-editing tools? Nat Rev Microbiol doi: 10.1038/nrmicro.2017.73.

Hutvagner G, Simard MJ. 2008. Argonaute proteins: key players in RNA silencing. Nat Rev Mol Cell Biol 9: 22–32.

Juliano C, Wang J, Lin H. 2011. Uniting Germline and Stem Cells: The Function of Piwi Proteins and the piRNA Pathway in Diverse Organisms. Annu Rev Genet 45: 447–469.

Jung SR, Kim E, Hwang W, Shin S, Song JJ, Hohng S. 2013. Dynamic anchoring of the 3’-end of the guide strand controls the target dissociation of Argonaute-guide complex. J Am Chem Soc 135: 16865–16871.

Katoh K, Standley DM. 2013. MAFFT Multiple Sequence Alignment Software Version 7: Improvements in Performance and Usability. Mol Biol Evol 30: 772–780.

Kaya E, Doxzen KW, Knoll KR, Wilson RC, Strutt SC, Kranzusch PJ, Doudna JA. 2016. A bacterial Argonaute with noncanonical guide RNA specificity. Proc Natl Acad Sci 113: 4057–4062.

Kinch LN, Ginalski K, Rychlewski L, Grishin NV. 2005. Identification of novel restriction endonuclease-like fold families among hypothetical proteins. Nucleic Acids Res 33: 3598–3605.

Koonin EV. 2017. Evolution of RNA- and DNA-guided antivirus defense systems in prokaryotes and eukaryotes: common ancestry vs convergence. Biol Direct 12. http://biologydirect.biomedcentral.com/articles/10.1186/s13062-017-0177-2 (Accessed March 1, 2017).

Kuramochi-Miyagawa S, Watanabe T, Gotoh K, Totoki Y, Toyoda A, Ikawa M, Asada N, Kojima K, Yamaguchi Y, Ijiri TW, et al. 2008. DNA methylation of retrotransposon genes is regulated by Piwi family members MILI and MIWI2 in murine fetal testes. Genes Dev 22: 908–917.

Laganeckas M, Margelevičius M, Venclovas Č. 2011. Identification of new homologs of PD-(D/E)XK nucleases by support vector machines trained on data derived from profile–profile alignments. Nucleic Acids Res 39: 1187–1196.

Lisitskaya L, Aravin AA, Kulbachinskiy A. 2018. RNA interference and beyond: structure and functions of prokaryotic Argonaute proteins. Nat Commun In review.

Liu J, Carmell MA, Rivas FV, Marsden CG, Thomson JM, Song J-J, Hammond SM, Joshua-Tor L, Hannon GJ. 2004. Argonaute2 Is the Catalytic Engine of Mammalian RNAi. Science 305: 1437–1441.

Liu Y, Esyunina D, Olovnikov I, Teplova M, Kulbachinskiy A, Aravin AA, Patel DJ. 2018. Accommodation of helical imperfections in Rhodobacter sphaeroides Argonaute ternary complexes with guide RNA and target DNA. Cell Rep 24: 1–10.

Ma J-B, Yuan Y-R, Meister G, Pei Y, Tuschl T, Patel DJ. 2005. Structural basis for 5’-end-specific recognition of guide RNA by the A. fulgidus Piwi protein. Nature 434: 666–70.

Makarova KS, Wolf YI, van der Oost J, Koonin EV. 2009. Prokaryotic homologs of Argonaute proteins are predicted to function as key components of a novel system of defense against mobile genetic elements. Biol Direct 4: 29.

Matsumoto N, Nishimasu H, Sakakibara K, Nishida KM, Hirano T, Ishitani R, Siomi H, Siomi MC, Nureki O. 2016. Crystal Structure of Silkworm PIWI-Clade Argonaute Siwi Bound to piRNA. Cell 167: 484–497 e9.

Miyoshi K. 2005. Slicer function of Drosophila Argonautes and its involvement in RISC formation. Genes Dev 19: 2837–2848.

Miyoshi T, Ito K, Murakami R, Uchiumi T. 2016. Structural basis for the recognition of guide RNA and target DNA heteroduplex by Argonaute. Nat Commun 7: ncomms11846.

Nakanishi K, Weinberg DE, Bartel DP, Patel DJ. 2012. Structure of yeast Argonaute with guide RNA. Nature 486: 368–374.

Olovnikov I, Chan K, Sachidanandam R, Newman DK, Aravin AA. 2013. Bacterial Argonaute Samples the Transcriptome to Identify Foreign DNA. Mol Cell 51: 594–605.

Park MS, Phan HD, Busch F, Hinckley SH, Brackbill JA, Wysocki VH, Nakanishi K. 2017. Human Argonaute3 has slicer activity. Nucleic Acids Res 45: 11867–11877.

Parker JS, Parizotto EA, Wang M, Roe SM, Barford D. 2009. Enhancement of the Seed-Target Recognition Step in RNA Silencing by a PIWI/MID Domain Protein. Mol Cell 33: 204–214.

Parker JS, Roe SM, Barford D. 2004. Crystal structure of a PIWI protein suggests mechanisms for siRNA recognition and slicer activity. EMBO J 23: 4727–4737.

Parker JS, Roe SM, Barford D. 2005. Structural insights into mRNA recognition from a PIWI domain–siRNA guide complex. Nature 434: 663–666.

Price MN, Dehal PS, Arkin AP. 2010. FastTree 2 – Approximately Maximum-Likelihood Trees for Large Alignments. PLOS ONE 5: e9490.

Rand TA, Ginalski K, Grishin NV, Wang X. 2004. Biochemical identification of Argonaute 2 as the sole protein required for RNA-induced silencing complex activity. Proc Natl Acad Sci U S A 101: 14385–14389.

Rehwinkel J, Behm-Ansmant I, Gatfield D, Izaurralde E. 2005. A crucial role for GW182 and the DCP1:DCP2 decapping complex in miRNA-mediated gene silencing. RNA 11: 1640–1647.

Remmert M, Biegert A, Hauser A, Söding J. 2012. HHblits: lightning-fast iterative protein sequence searching by HMM-HMM alignment. Nat Methods 9: 173.

Rivas FV, Tolia NH, Song J-J, Aragon JP, Liu J, Hannon GJ, Joshua-Tor L. 2005. Purified Argonaute2 and an siRNA form recombinant human RISC. Nat Struct Mol Biol 12: 340–349.

Schirle NT, Sheu-Gruttadauria J, MacRae IJ. 2014. Structural basis for microRNA targeting. Science 346: 608–613.

Sheng G, Zhao H, Wang J, Rao Y, Tian W, Swarts DC, van der Oost J, Patel DJ, Wang Y. 2014. Structure-based cleavage mechanism of Thermus thermophilus Argonaute DNA guide strand-mediated DNA target cleavage. Proc Natl Acad Sci 111: 652–657.

Sienski G, Dönertas D, Brennecke J. 2012. Transcriptional Silencing of Transposons by Piwi and Maelstrom and Its Impact on Chromatin State and Gene Expression. Cell 151: 964–980.

Soding J. 2005. Protein homology detection by HMM-HMM comparison. Bioinformatics 21: 951–960.

Song J-J, Smith SK, Hannon GJ, Joshua-Tor L. 2004. Crystal structure of Argonaute and its implications for RISC slicer activity. science 305: 1434–1437.

Swarts DC, Hegge JW, Hinojo I, Shiimori M, Ellis MA, Dumrongkulraksa J, Terns RM, Terns MP, van der Oost J. 2015a. Argonaute of the archaeon Pyrococcus furiosus is a DNA-guided nuclease that targets cognate DNA. Nucleic Acids Res 43: 5120–5129.

Swarts DC, Jore MM, Westra ER, Zhu Y, Janssen JH, Snijders AP, Wang Y, Patel DJ, Berenguer J, Brouns SJJ, et al. 2014a. DNA-guided DNA interference by a prokaryotic Argonaute. Nature 507: 258–261.

Swarts DC, Koehorst JJ, Westra ER, Schaap PJ, van der Oost J. 2015b. Effects of Argonaute on Gene Expression in Thermus thermophilus ed. L. Randau. PLOS ONE 10: e0124880.

Swarts DC, Makarova K, Wang Y, Nakanishi K, Ketting RF, Koonin EV, Patel DJ, van der Oost J. 2014b. The evolutionary journey of Argonaute proteins. Nat Struct Mol Biol 21: 743–753.

Verdel A, Jia S, Gerber S, Sugiyama T, Gygi S, Grewal SI, Moazed D. 2004. RNAi-mediated targeting of heterochromatin by the RITS complex. Science 303: 672–676.

Wang Y, Juranek S, Li H, Sheng G, Tuschl T, Patel DJ. 2008a. Structure of an argonaute silencing complex with a seed-containing guide DNA and target RNA duplex. Nature 456: 921–926.

Wang Y, Juranek S, Li H, Sheng G, Wardle GS, Tuschl T, Patel DJ. 2009. Nucleation, propagation and cleavage of target RNAs in Ago silencing complexes. Nature 461: 754–61.

Wang Y, Sheng G, Juranek S, Tuschl T, Patel DJ. 2008b. Structure of the guide-strand-containing argonaute silencing complex. Nature 456: 209–213.

Willkomm S, Makarova KS, Grohmann D. 2018. DNA silencing by prokaryotic Argonaute proteins adds a new layer of defense against invading nucleic acids. FEMS Microbiol Rev. https://academic.oup.com/femsre/advance-article/doi/10.1093/femsre/fuy010/4944904 (Accessed May 25, 2018).

Willkomm S, Oellig CA, Zander A, Restle T, Keegan R, Grohmann D, Schneider S. 2017. Structural and mechanistic insights into an archaeal DNA-guided Argonaute protein. Nat Microbiol 2: 17035–17035.

Yuan Y-R, Pei Y, Chen H-Y, Tuschl T, Patel DJ. 2006. A Potential Protein-RNA Recognition Event along the RISC-Loading Pathway from the Structure of A. aeolicus Argonaute with Externally Bound siRNA. Structure 14: 1557–1565.

Yuan Y-R, Pei Y, Ma J-B, Kuryavyi V, Zhadina M, Meister G, Chen H-Y, Dauter Z, Tuschl T, Patel DJ. 2005. Crystal Structure of A. aeolicus Argonaute, a Site-Specific DNA-Guided Endoribonuclease, Provides Insights into RISC-Mediated mRNA Cleavage. Mol Cell 19: 405–419.

Zander A, Holzmeister P, Klose D, Tinnefeld P, Grohmann D. 2014. Single-molecule FRET supports the two-state model of Argonaute action. RNA Biol 11: 45–56.

Zander A, Willkomm S, Ofer S, van Wolferen M, Egert L, Buchmeier S, Stockl S, Tinnefeld P, Schneider S, Klingl A, et al. 2017. Guide-independent DNA cleavage by archaeal Argonaute from Methanocaldococcus jannaschii. Nat Microbiol 2: 17034–17034.

